# Pattern-Induced Visual Discomfort and Its Temporal Summation Revealed by Pupillary Measures

**DOI:** 10.1101/2025.10.14.682064

**Authors:** Ron Meidan, Yoram S. Bonneh

## Abstract

Viewing repetitive striped patterns can induce pattern glare, experienced as visual discomfort (VD). While previous studies examined either pupillary responses or VD separately, few have investigated how they covary or evolve with repeated exposure. This study tested whether pupillary dynamics could serve as an objective “aversometer” — a physiological marker of individual visual sensitivity beyond subjective reports. Across four experiments (preliminary: n = 97; main: n = 70 for spatial frequency, n = 46 for central field size, n = 36 for central blank, with partial overlap), we manipulated spatial frequency, central field size, and surround field size of square-wave gratings (0.5–3 s) while measuring both discomfort and pupil size. Higher spatial frequencies and larger pattern areas elicited stronger pupillary constriction and greater discomfort, whereas repeated exposures produced cumulative increases in discomfort and decreases in baseline pupil size, consistent with visual strain rather than adaptation. To assess the potential of pupillometry as an aversometer, we examined individual differences in the main spatial-frequency experiment (controlled viewing distance, n = 42). A paradoxical pattern emerged: within participants, stronger stimuli produced greater constriction, but individuals with higher overall discomfort showed weaker constriction and stronger late redilation. Similar dissociations between subjective sensitivity and pupillary responses have been noted in studies of light-induced discomfort, suggesting that related mechanisms may contribute, although their specific physiological basis remains unclear. Overall, our findings clarify how pattern-induced discomfort evolves over time and across individuals and highlight pupillometry’s potential as a sensitive, objective tool for assessing visual sensitivity.

**Highlights:**

- Striped patterns systematically increased discomfort and pupillary constriction
- Repeated exposure led to progressive discomfort and shrinking baseline pupil
- Among high-sensitivity participants, weaker constriction and stronger redilation appeared
- The paradox may reflect interindividual autonomic differences under visual stress
- Pupillometry shows promise as an objective marker of visual sensitivity

## Introduction

Visual discomfort (VD) arises when viewing certain visual patterns, particularly high-contrast striped patterns with specific spatial frequencies (A. Wilkins et al., 1984). These patterns, resembling lines of text, can trigger symptoms including headaches, eyestrain, and perceptual distortions that may impair reading performance (A. J. Wilkins et al., 2007). The Pattern Glare Test quantifies susceptibility to visual stress using square-wave grating patterns, with the 3 cpd pattern typically eliciting the strongest visual distortions (Evans & Stevenson, 2008). Similarly, O’Hare and Hibbard (2011) further identified that spatial frequencies deviating from natural scenes produce maximal VD in most individuals. A systematic review by Evans and Allen (2016) defined visual stress as a distinct condition characterized by these symptoms when viewing repetitive patterns, likely resulting from hyperexcitability in the visual cortex (Monger et al., 2015).

While often discussed in the context of reading difficulties, similar visual sensitivity to striped patterns has been reported across several clinical populations. For example, individuals with conditions such as migraine (Hine & White, 2021; Shepherd et al., 2013) and autism (Torrens et al., 2024) might experience heightened sensitivity. Recent studies have also extended these findings to populations with chronic pain, as demonstrated by Ten Brink and Bultitude (2022). In many of these cases, current assessment methods rely primarily on subjective reports, which may be challenging to obtain from children or individuals with communication difficulties. Objective physiological measures may thus complement these assessments.

Pupillary responses offer one such measure, providing a potentially objective correlate of subjective discomfort, since the pupil is known to respond systematically to visual stimuli. (Barbur & Thomson, 1987) showed that pupil size varies with spatial frequency, while Ayzenberg et al. (2018) demonstrated that images of hole clusters that elicit aversive visual responses result in greater pupillary constriction. Lin et al. (2015) found that discomfort glare causes specific pupillary constriction responses that correlate with subjective ratings of discomfort.

Yet, while extensive research exists on pattern-induced visual discomfort and separate studies on pupillary responses to visual patterns, these two phenomena have rarely been examined simultaneously. The current study addresses this gap by investigating the relationship between VD and pupillary responses to striped patterns. We evaluated how pattern attributes affect both measures in parallel, aiming to connect subjective discomfort with an objective physiological response.

In the present study, we had three main aims. First, we sought to understand how specific pattern attributes (such as spatial frequency and field size) influence both visual discomfort and pupil size, and to establish a functional link between these two dependent variables. Second, we aimed to evaluate the feasibility of conducting such tests without standard chin rest conditions and explore the potential to develop a simplified, portable tool for assessing visual sensitivity. Third, we tested whether pupillary dynamics can potentially serve as an objective marker of visual sensitivity at the individual level.

## Materials and methods

### Participants

In the preliminary experiment, which consisted of two separate parts (objective and subjective), 97 adults participated in total (89 in the objective part, of whom 12 also completed the subjective part, plus 8 participants only in the subjective part). In Experiment 1 (spatial frequency), 70 adults participated. In Experiment 2 (central field size), 46 adults participated. In Experiment 3 (central blank), 36 adults participated (noting that certain participants took part in more than one of these main experiments). The overall gender distribution was nearly balanced (106 males, 111 females). Participant age was restricted to 18-35 years to minimize age-related variability in pupil size. All participants had normal or corrected-to-normal vision and were screened for any history of epilepsy or migraine. The experiments received approval from the Bar-Ilan University Institutional Review Board (IRB) Ethics Committee. Written informed consent was obtained from all participants, and all procedures adhered to IRB guidelines.

### Apparatus

Pupil size was recorded using a Tobii 4C eye tracker with a sampling rate of 90 Hz. The preliminary experiment utilized the tracker mounted on a 15.6-inch laptop monitor, while subsequent experiments employed a Yoga 7i laptop featuring a 14” FHD display (1920 x 1080). To ensure consistent display conditions, screen luminosity was set to maximum values, and automatic luminosity adjustments were disabled across all experiments. Stimulus presentation and data collection were managed through PSY, an in-house developed platform for psychophysical and eye-tracking experiments that has been validated in several of our previous studies (e.g, (Bonneh et al., 2015). No chin rest was used, to simulate field-deployable conditions suitable for diverse populations including children, with viewing distance continuously measured by the eye tracker. Exploratory analyses further examined subsets of participants based on their measured viewing distance, in order to assess whether accommodation or screen luminance at shorter distances might confound individual-level correlations. Although viewing distance was continuously tracked, it was only used in post-hoc analyses to identify individual differences (e.g., restricting participants to the 75–85 cm range) and was not enforced during data collection.The effective monitor visual angle was approximately 22.5º x 13.2º at the standard 75 cm viewing distance corresponding to the approximate average measured in the experiments. Eye tracking was conducted binocularly, with analyses focused on the left eye data. The visual discomfort (VD) rating scale ranged from 0 (no discomfort) to 5 (maximum discomfort), conceptually similar to established measures such as the Visual Discomfort Scale (Conlon & Hine, 2000) and the Leiden Visual Sensitivity Scale (Ten Brink et al., 2021). No explicit training was provided; instead, the first trial served as a familiarization trial for each participant.

### Stimuli and Procedure

#### Preliminary Experiment

Black and white square-wave gratings were presented at various spatial frequencies of 0.10, 0.33, 0.69, 1.64, and 7.20 cycles per degree. Luminance values were 0.08 cd/m^2^ for black stripes and 274 cd/m^2^ for the white background. Each stimulus was presented for 3000 ms followed by a 3000 ms white screen, with each condition repeated 5 times in random order. VD ratings and pupil measurements were conducted in two distinct sessions: an eye-tracking session where stimulus transitions were automatic, and a separate VD rating session where the next stimulus appeared only after the participant provided their rating for the current stimulus.

#### Experiment 1 - Spatial-Frequency

This experiment used square-wave gratings at spatial frequencies of 0.36, 0.47, 0.73, 1.72, and 4.97 cycles per degree presented on a brown background selected from preliminary testing as the least visually uncomfortable background color to reduce overall visual discomfort based on findings from a pilot study. The display area was kept constant at 21.10°×12.70°. Each stimulus was presented for 2500 ms followed by a minimum inter-stimulus interval of 1500 ms. The next stimulus was presented only after participants reported their VD rating by pressing buttons, resulting in an average time between trial onsets of approximately 5 seconds. Each condition was repeated 3 times in random order. Pupil size was continuously tracked throughout the experiment. Viewing distances varied within a range around the instructed distance of 75 cm, as estimated by the eye tracker.

#### Experiment 2 – Central Field Size

Black stripes (width: ^∼^0.15°, spacing: ^∼^ 0.1°) with a spatial frequency of ∼4 cycles per degree were presented at varying visual angles from 0.08°×0.08° to 12.40°×22.70°. All stimuli were displayed against a white background (274 cd/m^2^). Each stimulus was presented for 500 ms, followed by a minimum inter-stimulus interval of 1500 ms that was extended until participants provided their VD rating. Each condition was repeated 3 times in random order.

#### Experiment 3 – Surround Field Size

Stimuli consisted of a pattern with a spatial frequency of ^∼^ 4 cycles per degree extending over a total visual field of 21.10°×12.70°, with central blank windows of varying sizes (from 3.10°×3.10° to 12.10°×12.40°) presented against a white background. A control condition presented the full pattern without a blank window. Stimulus timing and rating procedure were identical to Experiment 2. Each condition was repeated three times in random order.

### Data Preprocessing

Pupil size data was analyzed to remove blinks and artifacts, following our previous (Kadosh et al., 2024). Blinks were detected when pupil size dropped to zero, and their exact onset and offset were refined using vertical eye movements exceeding 4 SD from baseline (calculated from the first third of the analysis window). We examined periods spanning 100 ms before and 150 ms after each blink. Blink windows outside the 250–750 ms range were marked as missing data, often due to head movement.

On average, approximately 25% of the data was missing due to blinks and tracking issues, which were partly due to free head and participant movement.

#### Epoch Extraction and Pupil Size Modulation

We extracted stimulus-aligned epochs spanning 0.5 s before onset to 3 s after offset for pupil size and eye position. To compute pupil size modulation (Figure 1), traces were normalized to the pre-onset average. Outliers (>2 SD per time point) were excluded within participants, and the data were then averaged across participants. Error bands represent ±1 standard error.

**Figure 1.**
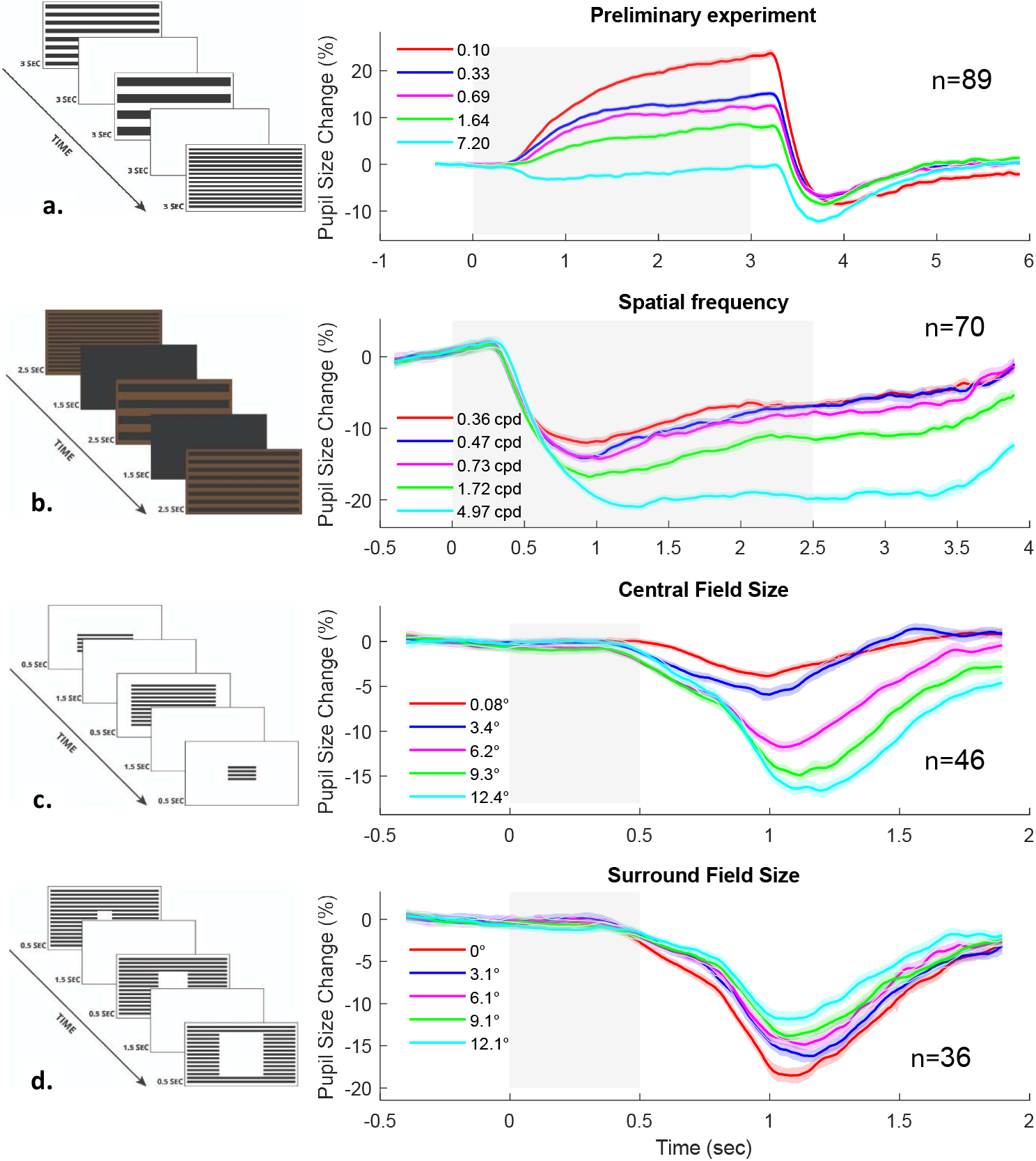
Pupillary responses to striped patterns across four experiments. Left: Square-wave gratings used in each experiment. Right: Corresponding pupillary time courses (gray shading indicates stimulus duration). (**a**) Preliminary experiment: Gratings with varying spatial frequencies (0.10–7.20 cpd) on a white background (3000 ms on, 3000 ms off), shown either automatically or with discomfort ratings. (**b**) Experiment 1: Similar gratings (0.36–4.97 cpd) on a brown-green background (2500 ms on, 1500 ms black screen) to reduce discomfort. (**c**) Experiment 2: Gratings with varying field sizes (1.0°–12.4°), shown for 500 ms, followed by a 1500 ms white screen. (**d**) Experiment 3: Gratings with different central blank sizes (0.1°–9.1°), same timing as (**c**). All experiments recorded pupil size and visual discomfort. Stripe luminance was constant (0.077 cd/m^2^); backgrounds were white (274.3 cd/m^2^) or brown (Exp. 1). The screen samples in a, b are similar to the actual stimulus with scaling, while the spatial-frequency shown in c, d was reduced to 30% for clarity.

#### Discrete Measures Extraction

For all experiments, we extracted baseline pupil size (pre-stimulus) and constriction responses. Baseline pupil size was defined as the average diameter during the 0.5 s preceding stimulus onset. Constriction was defined as the minimum pupil size (% change relative to baseline) within the 0.5–1.5 s window after stimulus onset, capturing the main constriction phase. In the preliminary experiment, constriction was instead measured in a 1–2.5 s window, to obtain initial estimates. In the main spatial-frequency experiment (Experiment 1), we additionally examined redilation, defined as the maximum pupil size (% change relative to baseline) in the 2–3 s post-stimulus recovery window.

#### Statistical Analysis

Linear trends were assessed using Linear Mixed Model (LMM) analysis (Hohenstein et al., 2017). Responses were fitted to a maximum likelihood model, with the above measures as dependent variables and stimulus properties or trial number as predictors. LMM p-values were computed at a 95% confidence level.

## Results

### The effect of pattern glare on pupil size and visual discomfort (VD) ratings

In the preliminary experiment, participants viewed striped gratings at different spatial frequencies. The study included 89 participants in the eye-tracking session and 20 in the separate subjective rating session, with 12 completing both sessions (n=97 unique participants). Pupil size was expressed as percentage change from baseline (Figure 1a). Because the background was white, the pupil dilated overall but also showed the Pupil Grating Response (PGR), a reflexive constriction linked to cortical processing of patterns that increased systematically with spatial frequency, consistent with previous reports (Barbur & Thomson, 1987; Hu et al., 2019). VD ratings, collected in a separate complementary session without eye tracking, also increased with spatial frequency, while showing the same trend as the pupillary responses even though they were measured in different sessions.

In the main experiments, pupil size and VD ratings were measured simultaneously. The experiments manipulated three stimulus attributes: spatial frequency (Experiment 1), central field size (Experiment 2), and surround field size (Experiment 3). As shown in Figure 1, pupil constriction varied with stimulus characteristics: greater constriction occurred with higher spatial frequency (Exp. 1, Fig. 1b), with larger central field size (Exp. 2, Fig. 1c), and with larger surround field size (Exp. 3, Fig. 1d; note reversed colors). Average viewing distances were 75.7 cm (SD=7.2), 72.3 cm (SD=9.9), and 76.3 cm (SD=12.3) for Experiments 1–3, respectively.

Figure 2 illustrates systematic relationships between stimulus variables and visual responses. In Figure 2a, pupillary constriction (blue, left axis) and VD ratings (orange, right axis) are plotted together across the three manipulations: spatial frequency (Exp. 1, n=70), central field size (Exp. 2, n=46), and central blank size (Exp. 3, n=36). Data were averaged within participants (with outlier exclusion, see Methods), then across participants, with ±1 SE error bars computed on demeaned data (with subject means subtracted to remove between-subject variability and better reveal within-subject effects of experimental manipulations).

**Figure 2.**
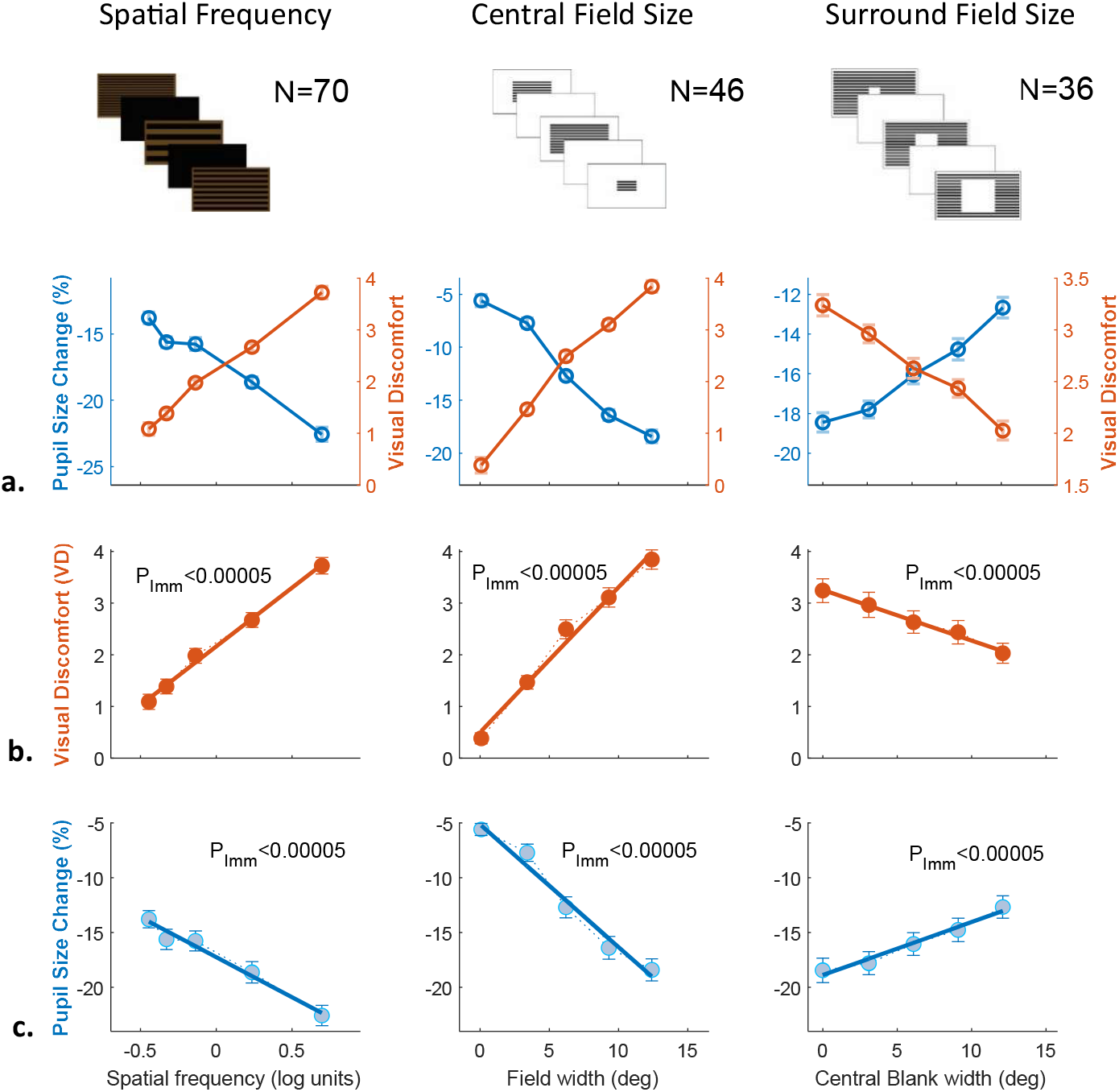
Systematic relationships between experimental variables and visual responses across three experimental paradigms. (**a**) Combined visualization of visual discomfort and pupillary responses. Visual discomfort (VD) ratings (orange circles, right y-axis) and pupillary constriction responses (blue circles, left y-axis) plotted against experimental variables. Left: Spatial frequency (Experiment 1, n=70), Middle: Central field size (Experiment 2, n=46), Right: Central blank size (Experiment 3, n=36). Both measures demonstrate systematic and opposite trends: as visual discomfort increases, pupillary constriction becomes stronger (more negative values). (**b**) Visual discomfort ratings plotted separately against each experimental variable. All relationships were statistically significant (P_lmm_< 0.00005), demonstrating systematic increases in subjective discomfort with increasing spatial frequency, field size, and decreasing central blank size. (**c**) Pupillary constriction responses plotted separately against each experimental variable. All relationships were statistically significant (P_lmm_< 0.00005), showing systematic changes in objective pupillary responses that mirror the subjective discomfort patterns. Data show means ± 1 SE.

Both measures showed monotonic, approximately linear relationships: pupil constriction increased (more negative change), whereas VD ratings increased with stimulus intensity. Figures 2b–c show each variable separately (VD and pupil constriction), using identical y-axis ranges for comparison across experiments. Significance was tested with linear mixed models (LMMs; see Methods). All six analyses (VD ratings and pupillary responses across Experiments 1–3) yielded p < 0.00005, confirming highly significant linear relationships. Analyses were based on discrete measures rather than continuous pupil time courses.

### The Relationship Between Visual Discomfort and Pupillary Measures

Following the demonstration that both VD and pupil constriction systematically vary with stimulus characteristics (Figure 2), we examined their direct correlation independent of specific stimulus parameters (Figure 3). To this end, we combined all stimulus conditions and plotted average pupil change (percentage from baseline) for each VD rating (0–5). This analysis allowed us to test whether subjective discomfort and pupillary responses are linked across different manipulations. The results revealed systematic relationships in all three experiments. Higher VD ratings were consistently associated with stronger pupil constriction (more negative change): spatial frequency (Exp. 1, p_lmm_<0.001), central field size (Exp. 2, p_lmm_ < 0.001), and central blank size (Exp. 3, p_lmm_ < 0.007). These findings demonstrate that subjective reports of visual discomfort are systematically associated with objective pupillary responses. Importantly, the correlations emerged when collapsing across conditions, showing that the link holds independently of specific stimulus attributes.

**Figure 3.**
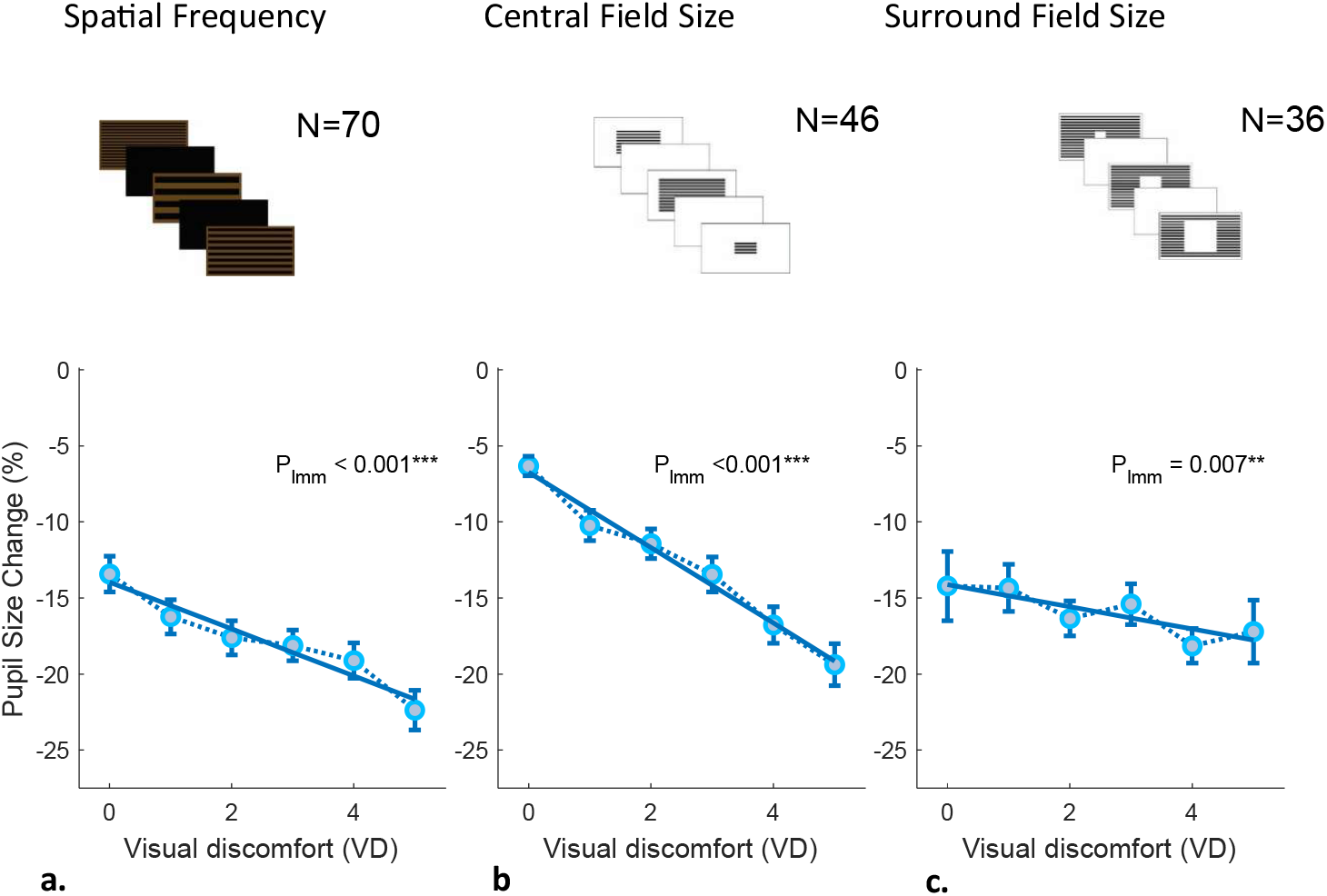
Relationships between visual discomfort ratings and pupillary measures across three experimental conditions. Graphs showing the relationship between visual discomfort (VD) ratings and pupil size change (%) across three experiments manipulating different stimulus properties: (**a**) Spatial frequency (70 participants); (**b**) Pattern field size (46 participants); (**c**) Central aperture size (36 participants). In all three conditions, increased visual discomfort was associated with greater pupil constriction (Spatial frequency: P_lmm_ <.001; Pattern field size: P_lmm_ <.001; Central aperture size: P_lmm_ =.0069). Each data point represents mean pupil response for participants who provided that specific VD rating level (0-5 scale). The number of participants contributing to each specific rating level varied (41-59 for spatial frequency, 29-43 for field size, 15-28 for aperture size) because participants provided repeated measures (rating multiple stimuli) and may not have used every possible rating level (0-5 scale). Linear mixed-effects models (LMM) accounted for repeated measures within participants.

### Temporal Summation of Visual Discomfort and Pupillary Responses

Figure 4 shows how VD ratings (left column) and pre-stimulus pupil size in mm (right column) changed across repeated exposures. The first trial in each experiment served as familiarization and was excluded from analysis. Across subsequent trials, VD ratings progressively increased in three of four experiments, reflecting an accumulation of visual discomfort with repeated stimulation. In the central field size experiment, significance was obtained only after excluding the two least aversive conditions (very small central fields, e.g., 0.08°), which consistently produced near-floor VD ratings. Removing these conditions allowed the analysis to focus on stimuli capable of eliciting measurable discomfort and revealed a clear cumulative effect (p = 0.0064 vs. 0.55; Fig. 4c). For the pupillary data, relative constriction (percentage change from baseline) did not show the expected temporal increase. This likely reflects the fact that baseline pupil size itself gradually shifted across trials, making relative measures less sensitive to cumulative changes. We therefore analyzed the pre-stimulus baseline pupil size in mm, which provided a clearer indicator of temporal effects. This analysis revealed significant trial-by-trial reductions in baseline pupil size across all experiments (p_lmm_ < 0.0005), paralleling the progressive increase in VD ratings.

**Figure 4.**
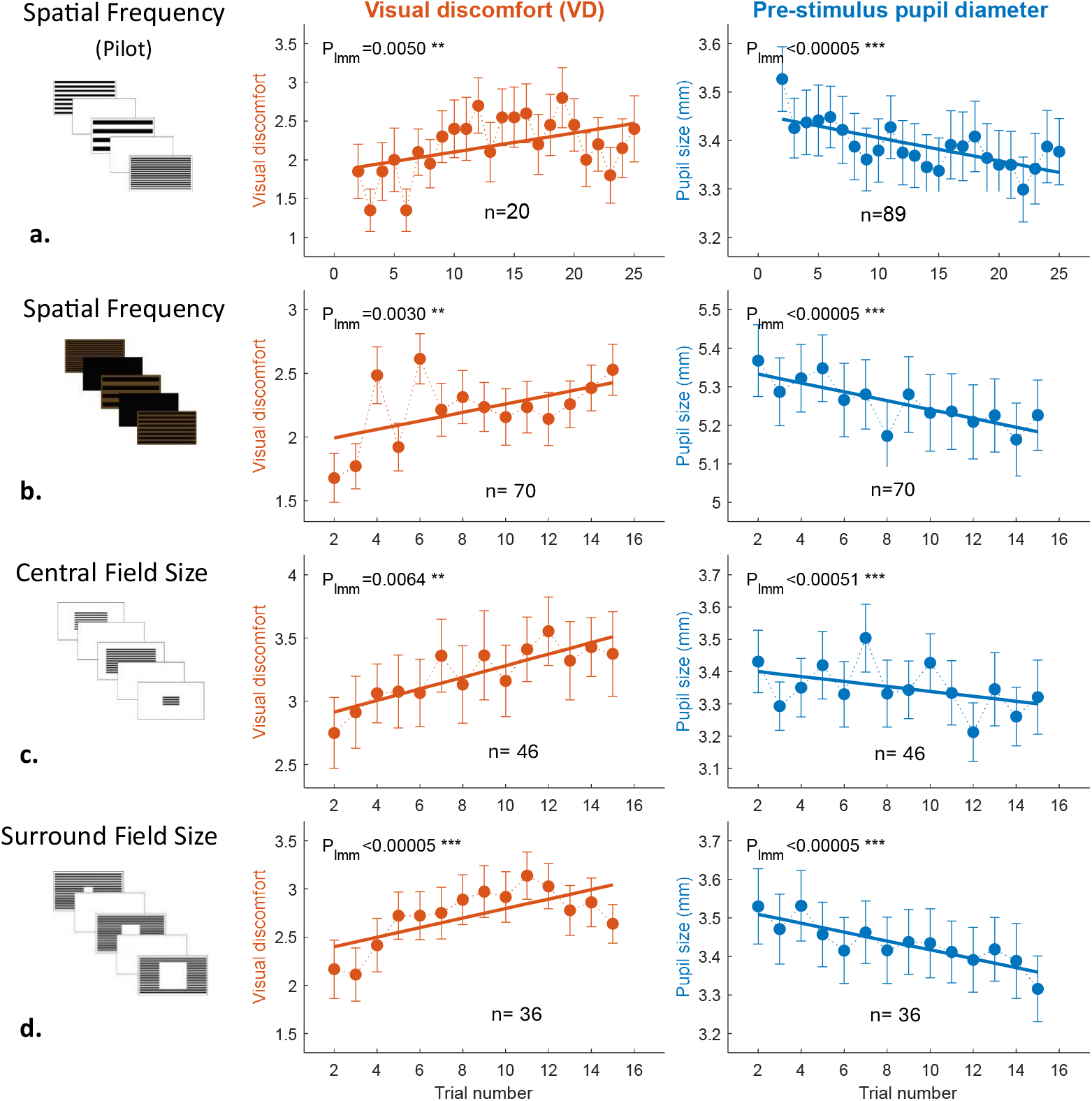
Temporal evolution of visual discomfort and pupil diameter across experiments. Visual discomfort (VD) ratings (left column, orange) and pre-stimulus pupil diameter in mm (right column, blue) as a function of trial number for: (**a**) preliminary spatial frequency with n=20 for VD and n=89 for pupil in separate sessions, (**b**) spatial frequency with n=70, (**c**) central field size with n=46, and (**d**) central aperture with n=36. The first trial in each experiment was excluded from analysis as it served to familiarize participants with the procedure. In central field size, the two smallest field conditions were excluded from analysis due to near-floor VD ratings. For pupil data, analyzed sample size per trial varied due to quality filtering: 75–85 in preliminary; 54–66 in spatial frequency; 22–32 in central field; 30–36 in central aperture. VD ratings increased across trials in three experiments; central field showed significance at p=0.0064. Pupil diameter decreased across trials in all experiments (p<0.0005). Error bars: SEM.

### Exploratory pupillary measures of individual differences in VD (“Aversometer”)

Both discomfort ratings and pupillary responses varied systematically with stimulus characteristics (Figure 2). To examine whether pupillary measures could capture individual differences in visual sensitivity, we conducted exploratory analyses in Experiment 1, which provided the largest sample size. A systematic scan of viewing distance ranges revealed that pupillary-discomfort correlations emerged only when participants maintained distances of 75–85 cm (n = 42 of 70). After outlier removal, we compared two groups: high-VD participants (average VD ≥ 2.5, n = 20) and low-VD participants (average VD < 2.5, n = 22). Both groups showed frequency-dependent patterns: VD ratings and peak constriction increased with spatial frequency (Fig. 5a−b). However, the groups differed markedly in their response profiles. The high-VD group reported consistently greater discomfort across all frequencies (Fig. 5A), yet exhibited weaker pupillary constriction (Fig. 5B) and stronger post-constriction recovery (Fig. 5C) compared to the low-VD group.

**Figure 5.**
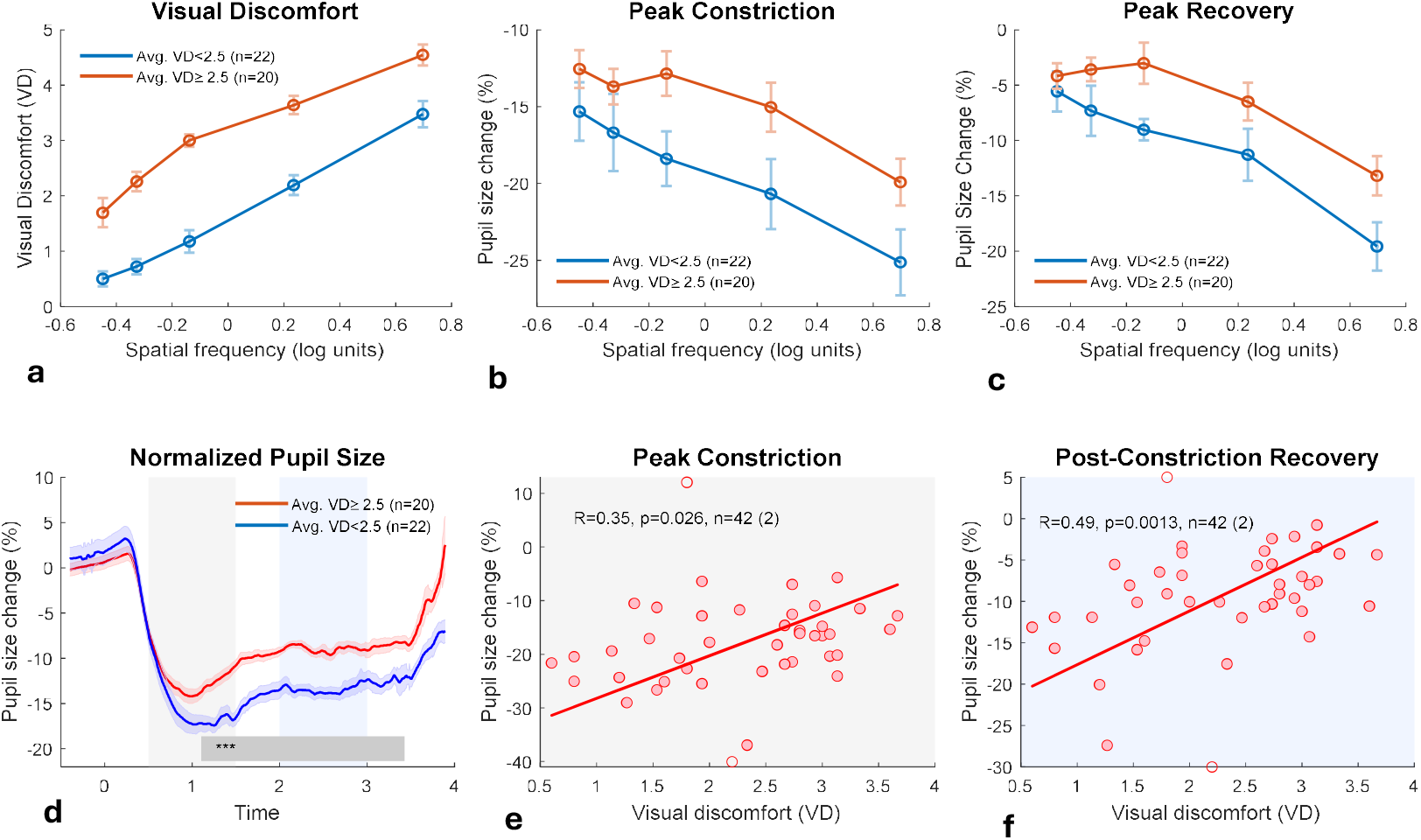
Pupillary responses and visual discomfort across spatial frequencies: group differences and individual correlations. (Top row) Group comparisons across five spatial frequencies [log_10_(freq) = −0.45 to 0.70, corresponding to 0.36-4.97 cycles per degree, cpd]. Participants were divided into high visual discomfort (Avg. VD ≥ 2.5, orange, n=20) and low visual discomfort (Avg. VD < 2.5, blue, n=22) groups based on average ratings. The number of participants per frequency bin varied slightly (18-20 per bin for both groups across measures) due to data quality filtering. (**a**) Visual discomfort ratings increase with spatial frequency in both groups, with the high-VD group consistently reporting greater discomfort. (**b**) Peak pupillary constriction shows frequency-dependent modulation, with stronger constriction (more negative values) at higher spatial frequencies in both groups. The high-VD group exhibits consistently weaker constriction responses across all frequencies. (**c**) Post-constriction pupillary recovery demonstrates frequency-dependent modulation, with weaker recovery (more negative values) at higher spatial frequencies. The high-VD group exhibits consistently stronger recovery (less negative values) across all frequencies. (Bottom row) Temporal dynamics and individual-level relationships. (**d**) Normalized pupillary response time courses (% change from pre-stimulus baseline). Gray shading (0.5−1.5 s) indicates the initial constriction phase; blue shading (2−3 s) indicates the recovery phase. Significant group differences emerge during the recovery period (1.1−3.4 s post-stimulus, p < 0.001, permutation test). (**e**) Individual-level correlation: higher average VD ratings are associated with weaker peak constriction (R = −0.35, p = 0.026, n = 42). (**f**) Individual-level correlation: higher average VD ratings are associated with stronger post-constriction recovery (R = 0.49, p = 0.0013, n = 42). Error bars represent standard error of the mean. All analyses restricted to viewing distance 75–85 cm. Statistical significance for time-series data assessed with permutation testing (1000 iterations). Outlier removal for scatter plots performed at 1.33 SD.

These group differences were most pronounced during the late recovery phase. Time-course analysis (Fig. 5D) revealed significant group divergence from 1.1–3.4 s post-stimulus (p < 0.001, permutation test), with high-VD participants showing faster pupil redilation. Individual-level correlations confirmed this pattern: higher average discomfort was associated with weaker peak constriction (R = –0.35, p = 0.026; Fig. 5E) and stronger recovery (R = 0.49, p = 0.0013; Fig. 5F). Overall, in Experiment 1, which manipulated spatial frequency, higher visual discomfort was consistently linked to weaker pupillary constriction and stronger post-constriction recovery, both in group comparisons and in individual-level correlations.

## Discussion

### Systematic Effects Across Experimental Manipulations

Our findings demonstrate robust stimulus–response relationships across all experimental manipulations. Both visual discomfort (VD) ratings and pupillary constriction showed highly significant correlations with pattern characteristics (all p < 0.00005) across experiments manipulating spatial frequency, central field size, and surround field size (Figure 2). The consistency of these relationships, despite variations in background colors and presentation durations, establishes spatial frequency and pattern area as fundamental drivers of both subjective and physiological responses to visual patterns.

Our findings are consistent with previous reports in the literature. Pupillary responses have been shown to vary systematically with the spatial frequency of grating patterns (Barbur & Thomson, 1987), often interpreted as a manifestation of the Pattern Glare Effect—where increased visual glare or higher *perceived* brightness leads to stronger pupillary constriction. This interpretation aligns with evidence that the pupillary response is modulated by perceived brightness rather than physical luminance alone, as demonstrated in studies using brightness illusions (Laeng & Endestad, 2012) and with images of natural light sources such as the sun (Binda et al., 2013). Together, these studies indicate that pupillary constriction can increase with higher spatial frequency or larger pattern area, even when physical luminance does not increase and may in fact be reduced. This matches our results: for example, in Experiment 2, larger pattern areas elicited stronger constriction despite containing less illuminated white area, whereas in the central blank window experiment (Experiment 3), larger windows emitted more light but led to weaker constriction and lower discomfort.

### Accumulation Rather Than Adaptation

Repeated exposures produced progressive increases in discomfort ratings (Figure 4, left) and decreases in pre-stimulus pupil size (Figure 4, right). This pattern represents a departure from typical visual adaptation, where responses usually weaken with repetition. For example, contrast adaptation in the visual system shows weakening over time (Ohzawa et al., 1982), which serves as a classic demonstration of visual adaptation. In contrast, our findings indicate cumulative strain, similar to temporal summation observed in pain research (Bar-Shalita & Cermak, 2015). Notably, our experiments did not include a separate familiarization session; instead, familiarization occurred during the very first trial, which likely explains why the temporal summation effect became statistically significant only after excluding trial 1. In the central field size experiment, excluding near-floor conditions revealed a robust effect (p shifting from 0.55 to 0.0064). This suggests that cumulative discomfort arises only when stimuli exceed a threshold of aversiveness. These results also have implications for experimental design. Repeated stimulus presentations, often used to increase statistical power, may themselves alter the responses being measured, as subjective discomfort and pupil size both accumulate over time.

### Individual Differences and the Sensitivity Paradox

At the group level, stimuli that produced higher discomfort ratings consistently elicited stronger pupillary constriction (Figure 2). However, across individuals, a counterintuitive pattern emerged: those reporting greater overall discomfort showed weaker stimulus-evoked constriction and stronger late-phase redilation (Figure 5). As shown in Figure 5b–f, high-sensitivity participants maintained larger baseline pupil diameters and more pronounced recovery. This dissociation between subjective and physiological measures represents a paradox of sensitivity, suggesting that individuals who experience stronger discomfort may exhibit altered autonomic balance rather than simply amplified responses.

The most direct clinical parallel to this pattern is found in migraine photophobia, a form of light-induced visual discomfort. Cortez et al. (2017) investigated the pupillary light response (PLR), a reflex to a pure light stimulus, and found that higher interictal photophobia was associated with a mixed autonomic dysfunction, specifically involving impairments in both pupillary constriction (parasympathetic mechanism) and the late redilation phase (sympathetic mechanism).

Our findings extend this principle to pattern-induced discomfort (Pattern Glare), a more complex visual stress that involves both luminance and high-spatial-frequency contrast. The observed pattern of weaker constriction and enhanced redilation in our high-discomfort group strongly mirrors the autonomic dysregulation reported in migraine, suggesting that the physiological basis for subjective visual sensitivity, whether triggered by pure light or structured patterns, may involve a common pathway of impaired autonomic regulation under visual load. This parallel supports the potential of pupillometry to serve as an objective marker for general visual hypersensitivity.

### Limitations

Several limitations should be acknowledged. First, we assumed that participants with higher discomfort ratings represent individuals with greater visual sensitivity, but this was not validated with standardized clinical measures. Future studies should combine pupillometry with validated sensitivity questionnaires and include clinical populations. Second, accommodation may have influenced responses. Participants who sat very close to the screen tended to report stronger discomfort and greater pupillary constriction, raising the possibility that accommodative effort contributed to these effects. However, within the 75–85 cm range accommodation effects were minimized, and no consistent relationship was found between viewing distance, pupil responses, or discomfort. Haigh et al. (Haigh et al., 2013) reported that uncomfortable patterns did not modulate accommodation, and suggested that weaker accommodation may be a consequence of visual discomfort or a coping mechanism rather than its cause. Xu et al. (Xu et al., 2015) showed that accommodative responses decrease as spatial frequency increases. Finally, we did not use a chinrest. This was intentional, to evaluate feasibility in populations such as children and to allow post-hoc filtering of viewing distances rather than enforcing rigid positioning. As a result, for the analysis of individual differences in visual discomfort, we excluded participants whose viewing distance frequently fell outside the reliable 75– 85 cm range, reducing the effective sample size for the data in Experiment 1 from n=70 to n=42. Nevertheless, within this controlled range, our results were consistent with prior literature such as Cortez et al. (2017).

## Conclusions

In summary, this study revealed three main findings. First, systematic effects were observed across all experimental manipulations, with both visual discomfort ratings and pupillary constriction increasing with higher spatial frequencies and larger pattern areas. Second, repeated exposures produced progressive increases in discomfort ratings and decreases in baseline pupil size, indicating accumulation of strain over time rather than typical adaptation. Finally, individual differences showed a paradoxical pattern: participants reporting the highest discomfort exhibited the weakest pupillary constriction.

### Declaration of Generative AI and AI-assisted Technologies in the Writing Process

During the preparation of this manuscript, the authors used Claude (Anthropic) to assist with language editing and formatting. After using this tool, the authors reviewed and edited all content as needed and take full responsibility for the content of the publication.

